# Decoding cell-type-specific alterations in Alzheimer’s disease through scRNA-seq and network analysis

**DOI:** 10.1101/2025.06.03.657612

**Authors:** Andrea Álvarez-Pérez, Lucía Prieto-Santamaría, Alejandro Rodríguez-González

## Abstract

Alzheimer’s Disease (AD) is a neurodegenerative disorder characterized by complex, cell-type-specific molecular alterations. This study integrates single-cell RNA sequencing (scRNA-seq) with network-based methodologies to decode transcriptional changes across major brain cell types in AD. Using scRNA-seq data from 432,555 single cells, we constructed Protein-Protein Interaction (PPI) networks specific to each cell type and assessed the differential expression of genes in diseased conditions. Our findings reveal that glutamatergic neurons and inhibitory interneurons exhibit the highest transcriptional dysregulation, while pericytes and endothelial cells show limited changes. The analysis identified significant enrichment of Differentially Expressed Genes (DEGs) within the AD protein module. Network analysis highlights highly connected proteins such as HSPB1, which is implicated in proteostasis, and CXCR4, which is involved in neuroinflammation. Our results underscore the importance of cell-type-specific approaches in AD research, demonstrating that neurons experience more extensive dysregulation, while vascular-associated cells play key roles in maintaining Blood-Brain Barrier (BBB) integrity. These insights emphasize the necessity of tailored therapeutic strategies addressing the heterogeneous molecular landscape of AD.

## I. Introduction

Neurological disorders are conditions of the central and peripheral nervous systems, including neurodegenerative diseases such as Alzheimer’s Disease (AD) and Parkinson’s Disease (PD), as well as developmental disorders like autism. These disorders are among the leading causes of disability worldwide, yet their etiology remains elusive due to phenotypic heterogeneity [1]. However, understanding neurological diseases faces several challenges. Cellular heterogeneity due to the complex nature of the brain tissue complicates the interpretation of transcriptomic changes, and correlation-based approaches often fail to establish causal relationships between genes and biological processes [2]. Furthermore, the selection of high-quality, well-curated datasets is crucial for generating reliable conclusions, particularly in multi-omics studies.

Single-cell RNA sequencing (scRNA-seq) has revolutionized disease research by enabling the identification of cell-type-specific transcriptional changes and high-resolution analysis of cellular heterogeneity and gene expression [3]. Unlike bulk RNA sequencing, scRNA-seq allows classification of cell subtypes, identification of activated gene networks, and analysis of intercellular interactions [4]. Studies in AD have revealed critical transcriptional alterations across cell types, such as differential regulation of the AD risk gene APOE in Oligodendrocyte Precursor Cells (OPCs) and astrocytes [5], and the disruptions in myelination-related processes across multiple cell types in the prefrontal cortex [6].

Network science and network medicine have emerged as transformative approaches for unraveling the molecular and systemic complexity of neurological disorders. In network medicine, relationships between nodes often represent direct physical interactions, or similarities in their interaction patterns with other molecules [7]. Within this approach, Protein-Protein Interaction (PPIs) networks serve as a powerful tool to explore functional modules associated with disease, as proteins with similar biological functions often interact closely [8].

Elucidating brain cell type specific gene expression and transcriptomic patterns is critical towards a better understanding of how cell-cell communications may influence brain functions and dysfunctions [9]. The integration of network-based methodologies and scRNA-seq has become a promising strategy for understanding the molecular underpinnings of neurological diseases, providing a system-wide perspective on disease mechanisms, revealing functionally relevant molecular alterations.

Several studies have applied such methodologies to investigate cell-type-specific changes in neurodegenerative disorders. Methods like SCPPIN integrate scRNA-seq data with PPIs to identify active molecular modules across different transcriptional states [8], while cell-type-specific PPI networks built from Differentially Expressed Genes (DEGs) in AD brains have revealed key pathogenic genes and disease-associated pathways, which may serve as potential therapeutic targets in AD [10]. Multiscale network modeling, such as BRETIGEA (BRain cEll Type specIfic Gene Expression Analysis), has shown the conservation of co-expression modules across cell types, further advancing our understanding of brain cellular organization and gene regulation [9].

Beyond PPI-based analyses, scRNA-seq studies and single-cell transcriptomic profiling of the superior frontal gyrus has revealed microglial gene expression alterations linked to AD risk genes, shedding light on cell-specific contributions to disease progression [11]. Additionally, integrative approaches combining snRNA-seq, snATAC-seq, and spatial transcriptomics have identified co-expression modules in multiple sclerosis, highlighting immune and glial cell interactions [12]. Computational tools like SCANet and conditional Cell-Specific Networks (c-CSNs), further enhance gene expression integration with network-based frameworks, improving cell clustering and disease modeling [13], [14].

The present study examines whether gene expression patterns in AD vary by cell type compared to healthy individuals. Using DEGs from scRNA-seq, we built cell-type-specific PPIs for eight major brain cell types. The findings help characterize AD-related molecular alterations through network-based analysis. The paper is organized as follows: Section II the materials and methods, including the data employed and the analyses performed. Section III details the results obtained and discusses them. And, finally, Section IV includes the conclusions and Section V identifies the limitations of the present work and the future research lines to be addressed.

## II. Materials and methods

### A. Materials

The dataset used in this study consists of scRNA-seq data from 432,555 single cells, publicly available in CellX-Gene^1^ and Synapse (syn52074156). It includes samples from AD (118,236 cells), frontotemporal dementia and progressive supranuclear palsy (215,525 cells), and normal controls (98,794 cells), with each cell expressing 29,968 genes.

Cells were collected from three brain regions—Broadmann area 4, the insular cortex, and the primary visual cortex—and classified into nine major cell types: astrocytes, glutamatergic neurons, oligodendrocyte precursor cells, oligodendrocytes, inhibitory interneurons, microglial cells, endothelial cells, pericytes, and T cells. Originally analyzed by Rexach et al. (2024), this dataset provides insights into selective neuronal vulnerability, disorder-specific glial-immune states, and gene regulatory networks in neurodegeneration [15].

All the files containing these data and the derived results are available through the following GitLab repository^2^.

### B. Methods

#### 1) Data preprocessing

The study focused on Alzheimer’s disease (AD) by selecting only the “normal” and “Alzheimer” conditions from the original dataset.

##### 1.1) Gene and cell filtering

Genes expressed in at least 10 cells within each cell type were retained. This ensures that only genes with sufficient expression are considered relevant for the analysis. In addition, cells expressing fewer than 200 genes were excluded from the analysis.

##### 1.2) Normalization and Log-transformation

Raw RNA-seq counts are dependent on sequencing depth. To address this, the total counts per cell were normalized to a constant value (10,000), enabling comparisons across cells with varying sequencing depths. A log-transformation (*log1p*) was applied to stabilize the variance and mitigate the impact of genes with extreme high abundance.

##### 1.3) Dataset splitting

The dataset was divided into 9 groups based on cell type. This division allows for the identification of DEGs between healthy and diseased conditions within each cell type, rather than in the combined dataset. From this point onward, analysis is conducted independently for each cell type.

#### 2) Differential expression analysis

Differential expression analysis was performed using the `sc.tl.rank_genes_groups()` function in Scanpy. This function identifies the genes that are most differentially expressed between the compared groups, using the “normal” condition as reference. The direction of the Log Fold Change (*logFC*) indicates whether a gene is up- or down-regulated in the group of interest, in our case, the diseased condition, with respect to the reference group.

Following these analyses, the cell type “T-cells” was excluded from further consideration due to the identification of only one DEG for this cell type.

#### 3) Characterization of AD module and integration with cell-type-specific DEGs

In the scRNA-seq dataset, genes are identified by ENSEMBL IDs, which are mapped to Entrez IDs, gene symbols, and Protein Accession Numbers using the NCBI API.

##### 3.1) AD module characterization

The AD module was characterized following the methodology described by Menche et al. (2015) [16], which involves identifying disease-associated proteins and the largest connected component (LCC) that they form in the PPI network. This subgraph, consisting of interconnected disease proteins, was statistically validated to ensure robustness. A detailed explanation is provided in Marín et al. (2024) [17].

##### 3.2) Integration of DEGs with the AD module

For each cell type, DEGs encoding proteins in the AD module were identified. We calculated the total number of DEGs and determined how many of them encoded known proteins. To assess their relevance to the AD module, we calculated the number of these proteins included within the module. Additionally, we computed the cell-type-specific sub-network module size and its statistical significance by selecting node sets with the same degree distribution as the seed genes, repeating this process 1,000 times. A z-score was calculated, followed by a p-value and an FDR-corrected adjusted p-value.

#### 4) Network analysis

##### 4.1) Network metrics calculation

To investigate the position of cell-type-specific DEG within the broader interactome, we calculated several network metrics, including degree, betweenness centrality, closeness centrality, and clustering coefficient. These metrics were computed across the entire interactome to understand how DEGs from each cell type are distributed within the network.

##### 4.2) Network analysis and visualization

To facilitate network visualization for the AD protein module, genes that were differentially expressed in at least one cell type were filtered from the full interactome, reducing the node count from 2,697 to 1,519. The AD disease module was then visualized using Gephi ^3^, where node size, based on degree, represented connectivity. This visualization helped highlight the relative importance of DEGs from different cell types within the AD module and their contribution to the network structure.

## III. Results and discussion

### 1) Characterization of Cell-Type-Specific Alterations in AD

To evaluate transcriptional differences across cell types in AD, we identified DEGs for each major brain cell type and evaluated their overlap with AD-associated protein modules. **Table I** summarizes the total DEGs per cell type, the subset of DEGs encoding proteins in the AD module, and the statistical significance of each cell-type-specific module.

**TABLE I:** CELL-TYPE-SPECIFIC DEGs AND MODULES.

| Cell type | Total DEGs | Cell type proteins in AD module | Cell type module size | adj p-value |
| --- | --- | --- | --- | --- |
| Glutamatergic neuron | 4436 | 733 | 277 | 7.68e-36 |
| Inhibitory interneuron | 4130 | 673 | 251 | 9.27e-29 |
| Oligodendrocyte | 3383 | 554 | 191 | 1.89e-20 |
| Astrocyte | 3176 | 532 | 183 | 8.56e-21 |
| Oligodendrocyte precursor cell | 1963 | 320 | 49 | 1.16e-81 |
| Microglial cell | 1585 | 334 | 62 | 4.73e-50 |
| Endothelial cell of vascular tree | 379 | 73 | 4 | 1.52e-18 |
| Pericyte | 111 | 20 | 1 | - |

The analysis reveals that glutamatergic neurons exhibit the highest number of DEGs (4,436), followed by inhibitory interneurons (4,130) and oligodendrocytes (3,383). Given that glutamate is the major excitatory neurotransmitter in the brain, the transcriptional dysregulation in glutamatergic neurons aligns with observed alterations in glutamate metabolism in familial AD patients [18]. Pericytes showed the fewest DEGs (111), with endothelial cells also exhibiting relatively small DEG counts. This may reflect the specialized roles of pericytes and endothelial cells within the neurovascular unit, regulating Blood-Brain Barrier (BBB) formation, maintenance, and vascular stability [19], resulting in less transcriptional dysregulation than cell types more directly affected by AD pathology.

Statistical validation of the cell-type-specific modules within the AD interactome showed significant enrichment for AD-associated proteins in multiple cell types, with the most substantial p-value observed in oligodendrocyte precursor cells (OPCs; adj. p = 1.16e-81).

### 2) Differentially expressed genes and shared molecular signatures across cell types

To identify common transcriptional alterations in AD, we analyzed the top 20 most up- and down-regulated DEGs across all cell types (**Fig. 1**).

**Fig. 1.**
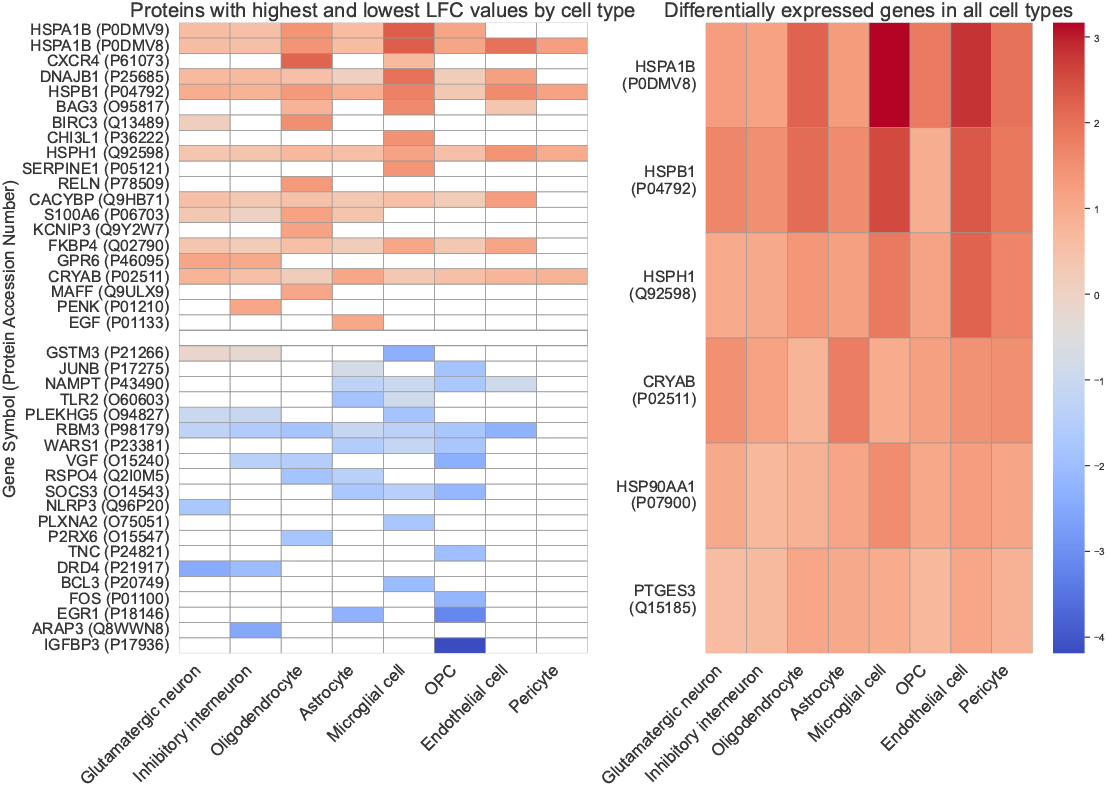
(Left) Heatmap displaying the top 20 most differentially up- and down-regulated genes and the proteins they encode. The color intensity represents the *logFC* values across cell types for Alzheimer’s disease relative to a healthy baseline condition. (Right) Heatmap showing the differentially expressed proteins present in all eight cell types.

Heatmap analysis revealed a consistent dysregulation of genes involved in protein folding, including HSPA1B, HSPB1, CRYAB, and HSP90AA1in all cell types, suggesting widespread perturbations in proteostasis mechanisms across the AD brain. Notably, HSPA1B was highly upregulated in microglial cells, endothelial cells, and oligodendrocytes, indicating a global cellular attempt to counteract oxidative stress and misfolded proteins. Microglia, as the brain’s primary immune cells, play a critical role in monitoring for signs of damage or infection in the brain [20]. In AD, the accumulation of misfolded tau and amyloid-beta triggers chronic microglial activation, leading to the upregulation of stress-response genes such as HSPA1B and DNAJB1 to mitigate protein aggregation.

Further analysis identified conserved dysregulated proteins, including CXCR4, a chemokine receptor critical for oligodendrocyte migration, maturation, and response to inflammatory signals. In AD patients, CXCR4 overexpression in oligodendrocytes may reflect an attempt to compensate for the loss of myelination by enhancing oligodendrocyte recruitment and survival in an inflammatory environment.

Additionally, IGFBP3 regulates insulin-like growth factor (IGF) signaling, which is crucial for OPC proliferation and differentiation. The pronounced negative differential expression in AD patients suggests that OPCs in the diseased brain experience impaired growth and differentiation, potentially contributing to defective remyelination and exacerbating neurodegeneration. While oligodendrocytes attempt to sustain myelin integrity, OPCs may be in an impaired differentiation state, limiting their ability to restock damaged oligodendrocytes and exacerbating myelin loss in the disease. Furthermore, endothelial cells and pericytes had fewer DEGs, likely reflecting their supportive roles in the neurovascular unit rather than direct involvement in neurodegeneration.

### 3) Representation of each cell type differential expression

The differential expression analysis identified key DEGs across the eight brain cell types in AD. **Fig. 2** displays volcano plots highlighting the five DEGs with the highest absolute *logFC* per cell type, where upregulated genes are marked in red and downregulated in blue. These DEGs were integrated into a PPI network of 1,519 proteins associated with AD module, generating eight cell-type-specific subnetworks. Node sizes were scaled by degree, and DEGs were colored by *logFC* to assess their positioning within the AD module.

**Fig. 2.**
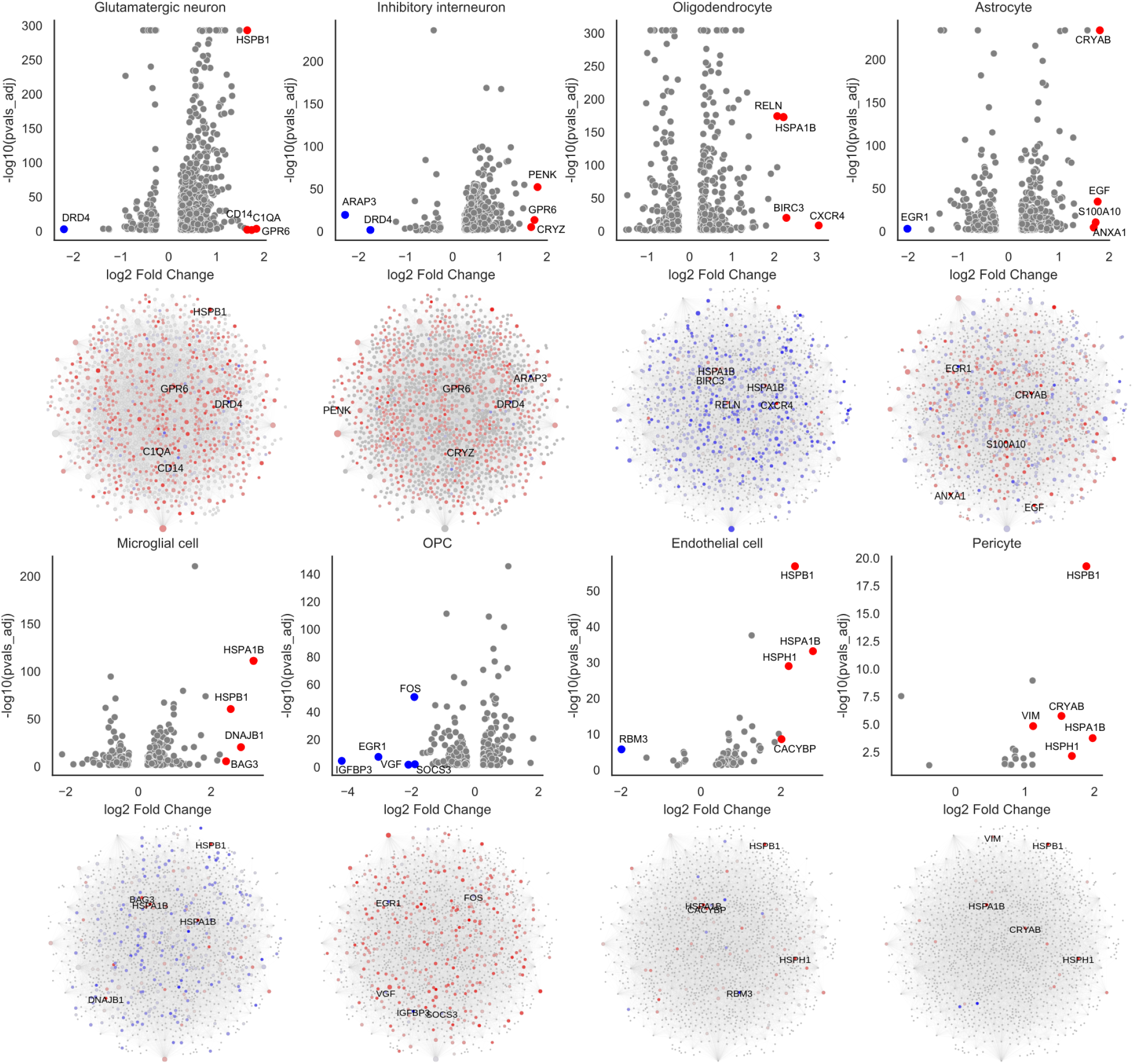
Volcano plots per cell type showing the DEGs *logFC* VS -log10(p-value adj), where top 5 with most absolute *logFC* value are highlighted, upregulated genes are marked in red and downregulated in blue. Below each volcano plot there is a PPI network representation of main AD module showing where node size is determmined by the degree, and the color by the magnitude and direction of *logFC*. Top 5 genes shown in the volcano plot are also annotated in the network.

Among the 28 unique genes annotated in the volcano plots, HSPB1 was the most highly connected node (degree = 320), appearing as a top DEG across multiple cell types. As a member of the heat shock protein family, its high connectivity suggests a crucial role in proteostasis, stress response, and neuroprotection.

VIM and FOS followed in connectivity (degree = 266 and 140, respectively). VIM, gene encoding vimentin, and upregulated in pericytes, may indicate a reactive response to BBB disruption, paralleling findings in pulmonary stress conditions, where increased VIM expression is associated with vascular dysfunction and cellular reactivity [21]. FOS, a transcription factor involved in neuronal plasticity and stress responses, and significantly downregulated in OPCs, may contribute to impaired differentiation and myelination, exacerbating AD-related white matter dysfunction [22].

Other highly connected DEGs, including BAG3, CACYBP, and ANXA1, exhibit cell-type-specific upregulation in microglia, endothelial cells, and astrocytes, respectively. BAG3 plays a key role in autophagy and protein quality control, particularly in microglia, which are essential for clearing mis-folded proteins; while ANXA1, an anti-inflammatory mediator whose expression in normal brain tissue is restricted to specific regions [23], is overexpressed in AD-associated astrocytes, reflecting a reactive response.

At the lower end of the AD PPI module, RELN and GPR6 exhibit the lowest connectivity (degree = 2). RELN downregulation in oligodendrocytes suggests altered neuronal homeostasis, while its absence in OPCs may indicate a differentiation block, as the main role of reelin is neurogenesis and neuronal polarization. GPR6 is a G protein-coupled receptor (GPR) involved in dopaminergic signaling, and its over expression in glutamatergic neurons in AD may affect synaptic modulation [24], though its low connectivity suggests localized rather than widespread effects. Similarly, VGF, ARAP3, and DRD4 (degrees < 5) had limited network influence, while other DEGs exhibited intermediate connectivity (degree = 10–100).

### 4) Network analysis of metrics distributions across cell types

To evaluate the network properties of DEGs within the AD interactome, we calculated degree, betweenness centrality, closeness centrality, and clustering coefficients for each cell-type-specific network. **Fig. 3** illustrates the log-transformed degree distribution across cell types. The degree of a node in a PPI network reflects its connectivity, providing insights into protein interactions within each cell type. The grey boxplot represents the general AD module, while blue boxplots indicate distributions in specific cell types. The “*n*” value represents the number of proteins in each subset.

**Fig. 3.**
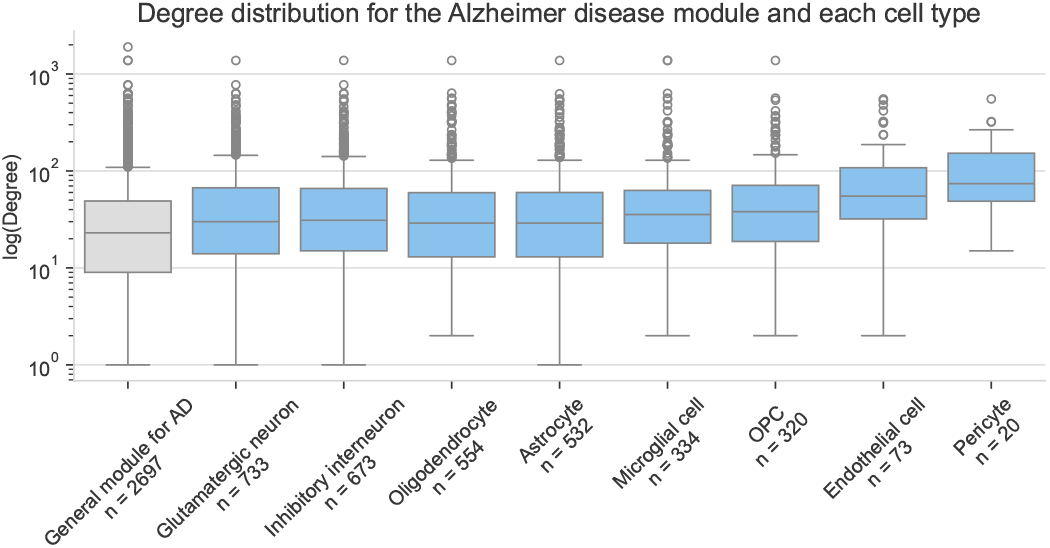
Degree distribution of DEGs within the general AD module, and stratified by cell type subnetworks.

The general AD module exhibits a lower median degree compared to individual cell types, likely due to its broader inclusion of proteins. Endothelial cells and pericytes show higher median degrees, suggesting stronger network integration related to BBB function, though differences in DEG numbers may affect this observation. Microglia and OPCs display high median degrees and broad distributions, indicating involvement of highly connected proteins probably reflecting its implication in neuroinflammation and remyelination. Oligodendrocytes and astrocytes exhibit substantial but slightly lower connectivity, likely reflecting their specialized roles in myelination and metabolic support. Glutamatergic and inhibitory interneurons share similar degree distributions, suggesting neuronal connectivity is influenced by synaptic and receptor-related proteins in the network.

## IV. Conclusions

This study investigated transcriptional alterations in different brain cell types and their association with AD, focusing on DEGs and their role in the AD-associated PPI network. The results showed significant dysregulation across various cell types, with glutamatergic neurons exhibiting the most extensive changes. Key alterations were observed in genes related to protein folding, stress responses, and neuroinflammation. A contrast between OPCs and mature oligodendrocytes indicated impaired differentiation in OPCs, potentially contributing to defective myelination in AD. The study also identified shared molecular signatures, such as up-regulated heat shock proteins and down-regulated IGFBP3 in OPCs, shedding light on cellular responses to stress and inflammation in AD. Network analysis highlighted hihly connected key proteins like HSPB1 and FOS as potential therapeutic targets. Cell type-specific sub-network metrics revealed unique differential expression patterns varying depending on the affected cell type.

Overall, our findings emphasize the complexity of AD pathogenesis, with diverse cellular mechanisms driving disease progression. Mapping differential gene expression across cell types and integrating this with the AD protein module provides valuable insights into how different brain cell types contribute to AD, offering potential insights for its characterization.

## V. Future steps and limitations

While this study offers valuable insights into cell-type-specific transcriptional alterations in AD, some limitations remain. The reliance on a single dataset may limit its applicability, as it should be validated using independent datasets. Future work should explore temporal dynamics using ATAC-seq data to identify early-stage alterations that precede AD pathology. Moreover, in addition to cell-type-specific classifications, constructing single-cell networks would provide deeper insights, as these networks could reveal unique cellular interactions and heterogeneity within each cell type, offering advantages over cell-type-based networks.

## Acknowledgments

The work is a result of the project “Data-driven drug repositioning applying graph neural networks (3DR-GNN)”, that is being developed under grant “PID2021-122659OB-I00” from the Spanish Ministerio de Ciencia e Innovación. A.A.-P. was granted by Universidad Politécnica de Madrid and Banco Santander for a predoctoral “Programa Propio” grant.

## Footnotes

1 https://cellxgene.cziscience.com/collections/c53573b2-eff4-4c5e-9ad0-b24d422dfd9b

2 https://medal.ctb.upm.es/internal/gitlab/disnet/network-medicine/network-medicine-and-single-cell-for-alzheimer

3 https://gephi.org/

## References

[1] W. M. Carroll, “The global burden of neurological disorders,” The Lancet Neurology, vol. 18, no. 5, pp. 418–419, May 2019. [Online]. Available: https://linkinghub.elsevier.com/retrieve/pii/S1474442219300298

[2] J.-C. García and R.-H. Bustos, “The genetic diagnosis of neurodegenerative diseases and therapeutic perspectives,” Brain Sciences, vol. 8, no. 12, p. 222, 12 2018.

[3] M. Ke, B. Elshenawy, H. Sheldon, A. Arora, and F. M. Buffa, “Single cell rna-sequencing: A powerful yet still challenging technology to study cellular heterogeneity,” BioEssays, vol. 44, no. 11, Sep. 2022. [Online]. Available: 10.1002/bies.202200084

[4] D. Huang, N. Ma, X. Li, Y. Gou, Y. Duan, B. Liu, J. Xia, X. Zhao, X. Wang, Q. Li, J. Rao, and X. Zhang, “Advances in single-cell rna sequencing and its applications in cancer research,” Journal of Hematology amp; Oncology, vol. 16, no. 1, Aug. 2023. [Online]. Available: 10.1186/s13045-023-01494-6

[5] A. Grubman, G. Chew, J. F. Ouyang, G. Sun, X. Y. Choo, C. McLean, R. K. Simmons, S. Buckberry, D. B. Vargas-Landin, D. Poppe, J. Pflueger, R. Lister, O. J. L. Rackham, E. Petretto, and J. M. Polo, “A single-cell atlas of entorhinal cortex from individuals with alzheimer’s disease reveals cell-type-specific gene expression regulation,” Nature Neuroscience, vol. 22, no. 12, p. 2087–2097, Nov. 2019. [Online]. Available: 10.1038/s41593-019-0539-4

[6] H. Mathys, J. Davila-Velderrain, Z. Peng, F. Gao, S. Mohammadi, J. Z. Young, M. Menon, L. He, F. Abdurrob, X. Jiang, A. J. Martorell, R. M. Ransohoff, B. P. Hafler, D. A. Bennett, M. Kellis, and L.-H. Tsai, “Single-cell transcriptomic analysis of alzheimer’s disease,” Nature, vol. 570, no. 7761, p. 332–337, May 2019. [Online]. Available: 10.1038/s41586-019-1195-2

[7] J. Barido-Sottani, S. D. Chapman, E. Kosman, and A. R. Mushegian, “Measuring similarity between gene interaction profiles,” BMC Bioinformatics, vol. 20, no. 1, p. 435, Dec. 2019. [Online]. Available: https://bmcbioinformatics.biomedcentral.com/articles/10.1186/s12859-019-3024-x

[8] F. Klimm, E. M. Toledo, T. Monfeuga, F. Zhang, C. M. Deane, and G. Reinert, “Functional module detection through integration of single-cell rna sequencing data with protein–protein interaction networks,” BMC Genomics, vol. 21, no. 1, Nov. 2020. [Online]. Available: 10.1186/s12864-020-07144-2

[9] A. T. McKenzie, M. Wang, M. E. Hauberg, J. F. Fullard, A. Kozlenkov, A. Keenan, Y. L. Hurd, S. Dracheva, P. Casaccia, P. Roussos, and B. Zhang, “Brain cell type specific gene expression and co-expression network architectures,” Scientific Reports, vol. 8, no. 1, Jun. 2018. [Online]. Available: 10.1038/s41598-018-27293-5

[10] X.-L. Wang and L. Li, “Cell type-specific potential pathogenic genes and functional pathways in alzheimer’s disease,” BMC Neurology, vol. 21, no. 1, Oct. 2021. [Online]. Available: 10.1186/s12883-021-02407-1

[11] Q. Wang, J. Antone, E. Alsop et al., “Single cell transcriptomes and multiscale networks from persons with and without alzheimer’s disease,” Nature Communications, vol. 15, p. 5815, 2024.

[12] M. L. Elkjaer, A. Hartebrodt, M. Oubounyt, A. Weber, L. Vitved, R. Reynolds, M. Thomassen, R. Rottger, J. Baumbach, and Z. Illes, “Single-cell multi-omics map of cell type–specific mechanistic drivers of multiple sclerosis lesions,” Neurology Neuroimmunology & Neuroinflammation, vol. 11, no. 3, May 2024. [Online]. Available: 10.1212/NXI.0000000000200213

[13] M. Oubounyt, L. Adlung, F. Patroni, N. K. Wenke, A. Maier, M. Hartung, J. Baumbach, and M. L. Elkjaer, “Inference of differential key regulatory networks and mechanistic drug repurposing candidates from scrna-seq data with scanet,” Bioinformatics, vol. 39, no. 11, Oct. 2023. [Online]. Available: 10.1093/bioinformatics/btad644

[14] L. Li, H. Dai, Z. Fang, and L. Chen, “c-csn: Single-cell rna sequencing data analysis by conditional cell-specific network,” Genomics, Proteomics & Bioinformatics, vol. 19, no. 2, p. 319–329, Mar. 2021. [Online]. Available: 10.1016/j.gpb.2020.05.005

[15] J. E. Rexach, Y. Cheng, L. Chen, D. Polioudakis, L.-C. Lin, V. Mitri, A. Elkins, X. Han, M. Yamakawa, A. Yin, D. Calini, R. Kawaguchi, J. Ou, J. Huang, C. Williams, J. Robinson, S. E. Gaus, S. Spina, E. B. Lee, L. T. Grinberg, H. Vinters, J. Q. Trojanowski, W. W. Seeley, D. Malhotra, and D. H. Geschwind, “Cross-disorder and disease-specific pathways in dementia revealed by single-cell genomics,” Cell, vol. 187, no. 20, pp. 5753–5774.e28, Oct. 2024. [Online]. Available: 10.1016/j.cell.2024.08.019

[16] J. Menche, A. Sharma, M. Kitsak, S. Ghiassian, M. Vidal, J. Loscalzo, and A.-L. Barabási, “Uncovering disease-disease relationships through the incomplete human interactome,” Science (New York, N.Y.), vol. 347, no. 6224, p. 1257601, Feb. 2015. [Online]. Available: https://www.ncbi.nlm.nih.gov/pmc/articles/PMC4435741/

[17] M. M. Tercero, L. Prieto-Santamaría, and A. Rodríguez-González, “Exploring drug repurposing opportunities for schizophrenia: A network medicine approach,” in 2024 IEEE 37th International Symposium on Computer-Based Medical Systems (CBMS). IEEE, Jun. 2024, p. 106–111. [Online]. Available: 10.1109/CBMS61543.2024.00026

[18] L. Mosconi, A. Pupi, and M. J. De Leon, “Brain glucose hypometabolism and oxidative stress in preclinical alzheimer’s disease,” Annals of the New York Academy of Sciences, vol. 1147, no. 1, p. 180–195, Dec. 2008. [Online]. Available: 10.1196/annals.1427.007

[19] E. A. Winkler, A. P. Sagare, and B. V. Zlokovic, “The pericyte: A forgotten cell type with important implications for alzheimer’s disease?” Brain Pathology, vol. 24, no. 4, p. 371–386, Jun. 2014. [Online]. Available: 10.1111/bpa.12152

[20] J. Miao, H. Ma, Y. Yang, Y. Liao, C. Lin, J. Zheng, M. Yu, and J. Lan, “Microglia in alzheimer’s disease: pathogenesis, mechanisms, and therapeutic potentials,” Frontiers in Aging Neuroscience, vol. 15, Jun. 2023. [Online]. Available: 10.3389/fnagi.2023.1201982

[21] T. Klouda, Y. Kim, S.-H. Baek, M. Bhaumik, Y. Li, Y. Liu, J. C. Wu, B. A. Raby, V. d. J. Perez, and K. Yuan, “Specialized pericyte subtypes in the pulmonary capillaries,” The EMBO Journal, vol. 44, no. 4, p. 1074–1106, Jan. 2025. [Online]. Available: 10.1038/s44318-024-00349-1

[22] K. G. Johnston, B. T. Berackey, K. M. Tran, A. Gelber, Z. Yu, G. R. MacGregor, E. A. Mukamel, Z. Tan, K. N. Green, and X. Xu, “Single-cell spatial transcriptomics reveals distinct patterns of dysregulation in non-neuronal and neuronal cells induced by the trem2r47h alzheimer’s risk gene mutation,” Molecular Psychiatry, vol. 30, no. 2, p. 461–477, Aug. 2024. [Online]. Available: 10.1038/s41380-024-02651-0

[23] Z. B. White, S. Nair, and M. Bredel, “The role of annexins in central nervous system development and disease,” Journal of Molecular Medicine, vol. 102, no. 6, p. 751–760, Apr. 2024. [Online]. Available: 10.1007/s00109-024-02443-7

[24] M. S. Alavi, A. Shamsizadeh, H. Azhdari-Zarmehri, and A. Roohbakhsh, “Orphan g protein-coupled receptors: The role in cns disorders,” Biomedicine and; Pharmacotherapy, vol. 98, p. 222–232, Feb. 2018. [Online]. Available: 10.1016/j.biopha.2017.12.056

